# Representation learning of RNA velocity reveals robust cell transitions

**DOI:** 10.1101/2021.03.19.436127

**Authors:** Chen Qiao, Yuanhua Huang

**Affiliations:** School of Biomedical Sciences, University of Hong Kong, Hong Kong SAR; School of Biomedical Sciences & Department of Statistics and Actuarial Science, University of Hong Kong, Hong Kong SAR

**Keywords:** Single-cell RNA velocity, Autoencoder, Cellular transitions

## Abstract

RNA velocity is a promising technique to reveal transient cellular dynamics among a heterogeneous cell population and quantify their transitions from single-cell transcriptome experiments. However, the cell transitions estimated from high dimensional RNA velocity are often unstable or inaccurate, partly due to the high technical noise and less informative projection. Here, we present VeloAE, a tailored representation learning method to learn a low-dimensional representation of RNA velocity on which cell transitions can be robustly estimated. From various experimental datasets, we show that VeloAE can both accurately identify stimulation dynamics in time-series designs and effectively capture the expected cellular differentiation in different biological systems. VeloAE therefore enhances the usefulness of RNA velocity for studying a wide range of biological processes.

## 1 Introduction

Single-cell RNA sequencing (scRNA-seq), by probing the full transcriptome of many individual cells, has become a revolutionary tool to study the cellular dynamic processes, such as cell cycle, cell differentiation, and organ genesis [1]. Numerous computational methods have been developed for trajectory inference over the past few years [2], covering various modern techniques, including principled graph fitting [3], diffusion map [4], Gaussian processes [5], optimal transport [6]. However, this task is still highly challenging, especially for automatically identifying the directionality of the inferred trajectory. A fundamental reason is that all these methods can only access to the current-state transcriptomes, but lack indications of past and / or future states.

Besides the conventional use of mature mRNAs, most scRNA protocols, including poly(A) enriched ones, contain a non-negligible proportion of nascent RNAs, reflecting its intrinsic RNA dynamics [7, 8]. Such informative unspliced RNAs have further been formulated into a novel concept of RNA velocity, i.e., the time derivative of mature mRNAs, hence indicating the past and future states of each gene and consequently the transition directionality between cells [8]. This technique has been further extended to a full dynamical model with a probabilistic setting [9]. However, despite its increasing applications and wide success, RNA velocities are often found to be unrobust or inaccurate for identifying the cellular transitions [10]. This is partly due to the nature of low contents of unspliced RNAs and the high noise in scRNA-seq data, and partly due to the lack of effective way for integrating the high transcriptome dimensions when projecting the cellular states. For the latter, one possible solution is to identify a set of dynamic related genes, e.g., by manual curation [10] or by prioritization with supervised covariates [11]. Instead of selecting a sub set of genes, here we proposed veloAE, a tailored representation learning method, to address the above challenges by constructing a joint lower-dimensional representation of both current mRNA states and their velocity vector. Hence, the cell transitions can be estimated in the lower-dimensional instead of original space. This framework is not only highly capable of de-noising the scRNA-seq count data [12], but also leverages its fully unsupervised manner to automatically learn an informative representation whereby cellular transitions can be robustly estimated.

## 2 Results

### 2.1 High level description of veloAE

Briefly, veloAE is a principled autoencoder, a common choice of representation learning in the form of neural network [13]. In this framework, the encoder sequentially consists of a conventional encoder, i.e., multi-layer perceptrons (MLP) with a single hidden layer, and a graph convolutional network (GCN) module (i.e., cohort aggregation in Fig. 1; Methods). The GCN aims to smooth the pre-encoded latent vector between neighbour cells by leveraging a predefined adjacent graph based on transcriptome similarity, e.g., K-nearest neighbour graph by default. For the decoder, we introduce an attentive combination module, which has been recently proposed for machine translation and soon popularized for a wide range of tasks [14, 15], so as to strengthen the biological interpretation of the latent dimensions here (Methods).

**Figure 1.**
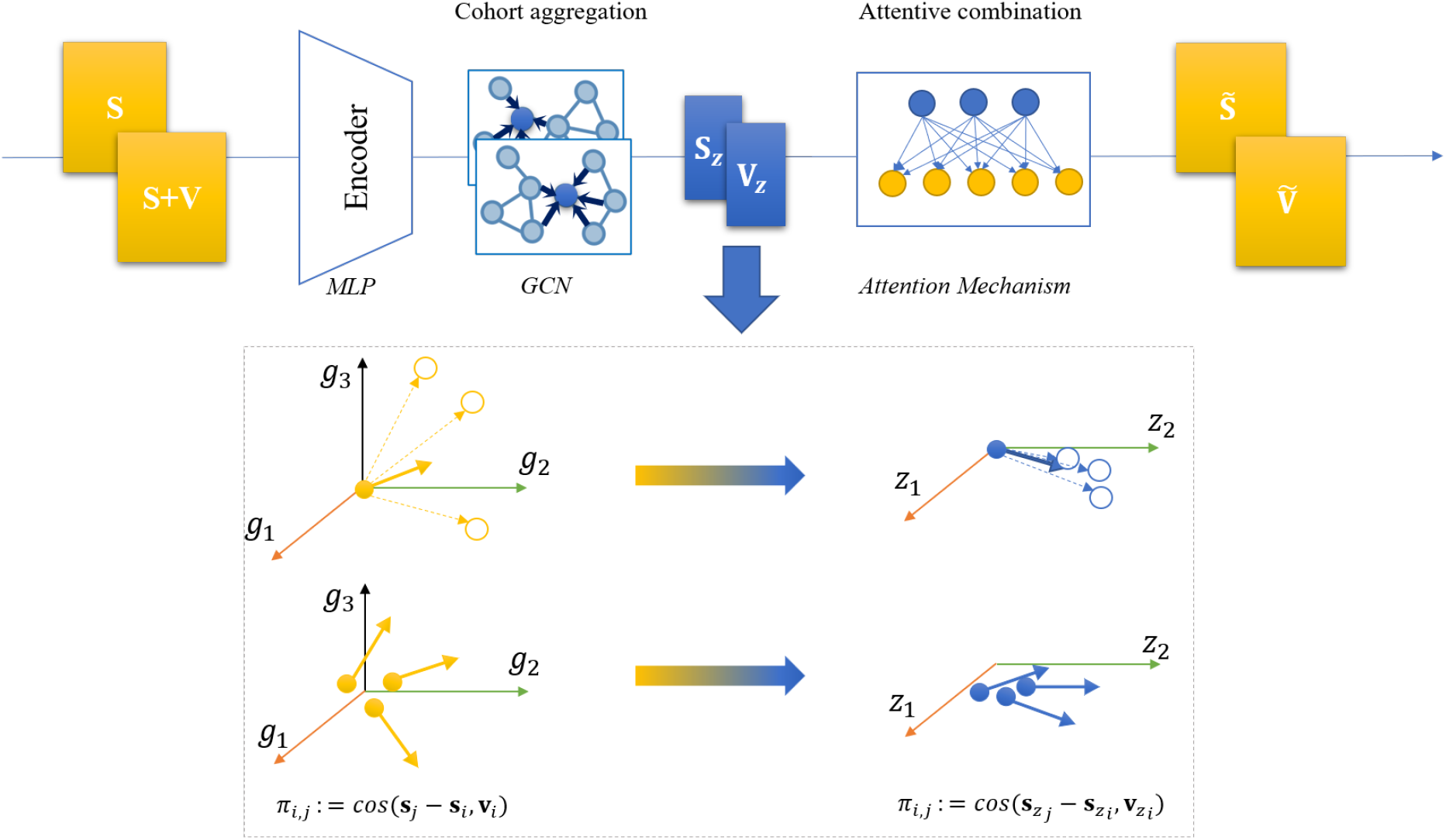
Overview of the veloAE with exemplified low-dimensional projection effects. Compared to standard autoencoder, veloAE has a cohort aggregation module with a graph convolutional network (GCN) in the encoder and an attentive combination module as decoder. Examples in box: the cell transition probability *π*_*i,j*_ and its directionality are smoothed and corrected from original space (left) to a learned low-dimensional space (right) for one cell to a multi-cell state (top) or multiple cells to a common differentiation direction (bottom).

The veloAE model will be jointly fit to the observed cell-by-gene count matrices respectively for spliced RNAs, unspliced RNAs, and optionally for aggregated expression if not redundant. Then both the velocity ***v*** and spliced RNAs ***s*** can be encoded to the same lower-dimensional representations (i.e., ***v***_***z***_ and ***s***_***z***_) by this fitted model (Methods; Fig. 1). Therefore, the cell transition probabilities can be computed in the lower-dimensional space based on ***v***_***z***_ and ***s***_***z***_, which is expected to both de-noise the data and to identify more informative projections (see examples in Fig. 1, and Methods section 4.2). Thanks to leveraging the computing power of graphic processing units (GPUs), veloAE can be fitted within around 7.2 minutes (20000 epochs) for around 3300 cells (see details of dataset statistics in Methods).

Moreover, for quantitative evaluation of velocity estimation, we additionally propose two metrics: Cross-Boundary Direction Correctness (CBDir) and In-Cluster Coherence (ICVCoh), for scoring the direction correctness and coherence of estimated velocities (see Section 4.5 for details). These metrics can complement the usual vague evaluation with mainly visual plotting of velocity filed.

### 2.2 veloAE corrects cell transitions in time-series stimulations

In order to evaluate the performance of veloAE in cell transition identification, we first applied it to a proof-of-principle design, where neurons are stimulated with KCl for 0, 15, 30, 60 and 120 minutes [10], hence the cellular transcriptomes are expected to transit along the dense stimulation time. In order to assess if RNA velocity is able to reconstruct the expected cell transitions, we estimated it from intronic and exonic reads by scVelo [9] (see Methods). However, we found the default RNA velocity in the original gene space struggles to identify the correct transition directions in the early time points for both scVelo’s stochastic mode (Fig. 2a) and its dynamical setting (Supplementary Fig. S1). We therefore ask whether most genes have the velocity at 0-min cells direct to the stimulated cells, by measuring the median velocity for 0-min cells (i.e., 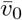) and their median expression differences to 15-min cells (i.e., 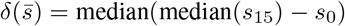. Surprisingly, the relation between velocity 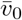 and forward expression difference 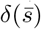 are largely stochastic (Pearson’s R=−0.042), and a substantial fraction of genes (50.7%) have opposite signs between them, indicating a backward transition (Fig. 2c). Similar pattern is also found in scVelo’s dynamical mode (Supp. Fig. S1). In other words, contradict transition directions widely exist among different genes (for example Arih1 has forward direction while Cltc has backward direction; Fig. 2c and Supp. Fig. S3), which hinders the identification of the expected direction in early time points.

**Figure 2.**
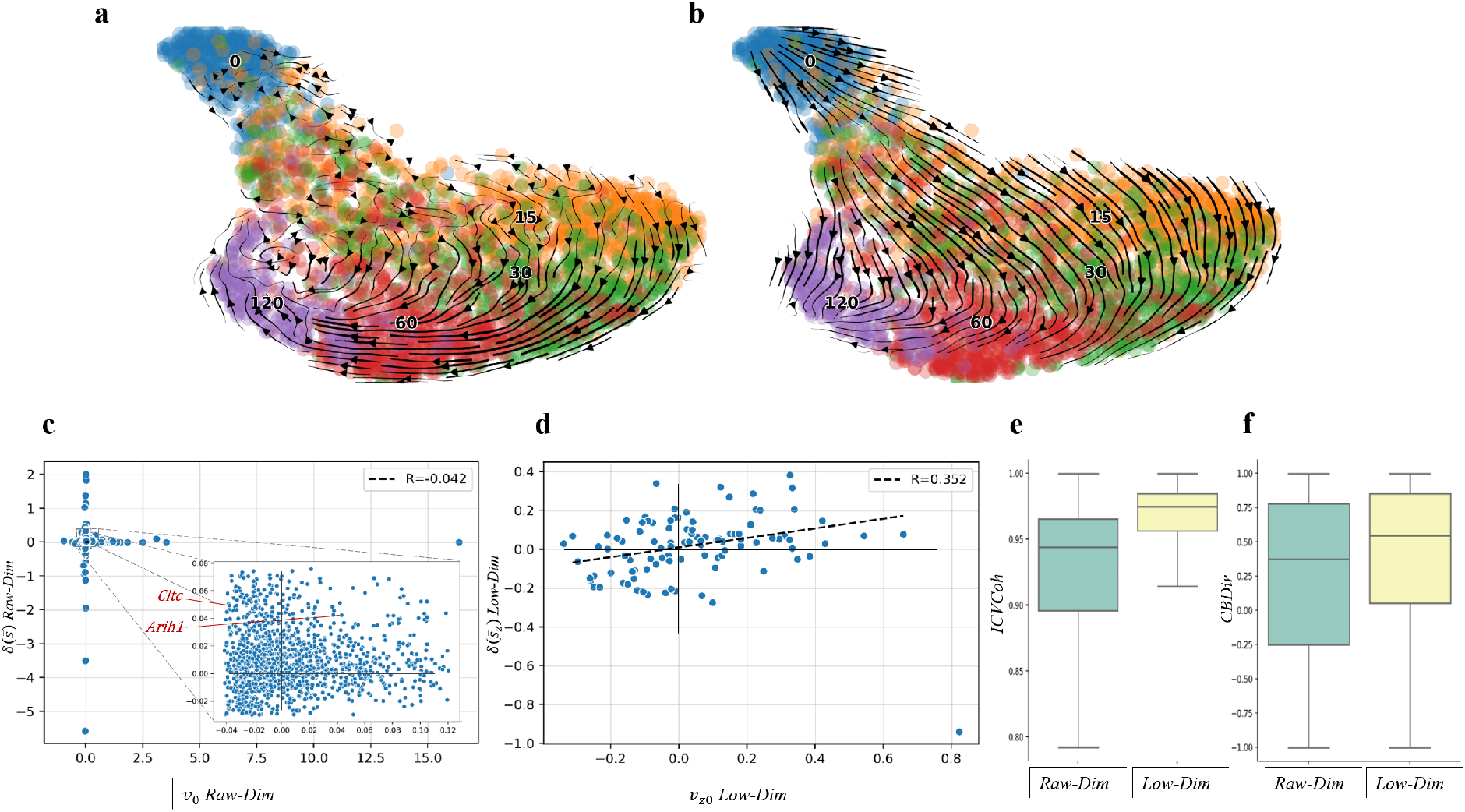
Results and Analysis on scNTseq. **a**, scVelo Stochastic Mode in Raw Gene Space; **b**, Velocity Projected into Low-Dimensional Space; **c**, Scatter plot of 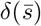 (15-0) over 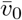 in the Raw Space; **d**, Scatter plot of 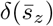 (15-0) over 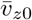 in the Low-Dim Space; **e**, In-cluster Coherence Scores; **f**, Cross-Boundary Direction Correctness (A->B) Scores.

On contrast, the learned representation in a 100-dimensional space by veloAE can largely retain a positive relation between 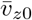 and 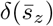 (Pearson’s R=0.352 on 98% interval) and more consistent signs between them (66.0%, Fig. 2d). Therefore, the cell transitions estimated from these latent dimensions are remarkably corrected, especially with a clear direction between 0 min to 15min (Fig. 2b). Overall, the veloAE’s representation not only substantially enhances the proportion of correct direction between two proximate states (mean CBDir: 0.253 by scVelo vs 0.402 by veloAE; Fig. 2f; Methods), but also increases the transition coherence within each sub group of cells (mean ICVCoh from 0.914 by scVelo to 0.964 by veloAE; Fig, 2e; Methods).

### 2.3 veloAE strengthens directionality in oligodendrocyte lineages

Next, we applied veloAE to a well studied data set on neuron genesis, where the development of mouse dentate gyrus was measured at two time points (P12 and P35) with scRNA-seq data (10x Genomics) [9, 16]. As demonstrated in the scVelo paper [9], major differentiation lineages, e.g., neuroblasts developing into granule cells, can be successfully identified by its both stochastic and dynamical modes. However, sub-lineages remain challenging to be identified, particularly, the differentiation from oligodendrocyte precursor cells (OPCs) into myelinating oligodendrocytes (OLs). Only the dynamical mode is able to moderately detect the right direction between them [9], and stochastic mode rather returns erroneous transitions (Fig. 3a). This challenge occurs partly due to that both cell types are in sub-stationary states and transient cells are limited. By visualizing the consistency between velocities at OPCs and its expression difference to OLs, we found there is only weak relation between the velocities and the expression difference (Pearson’s R=0.093) and 41.1% genes show opposite signs between them.

**Figure 3.**
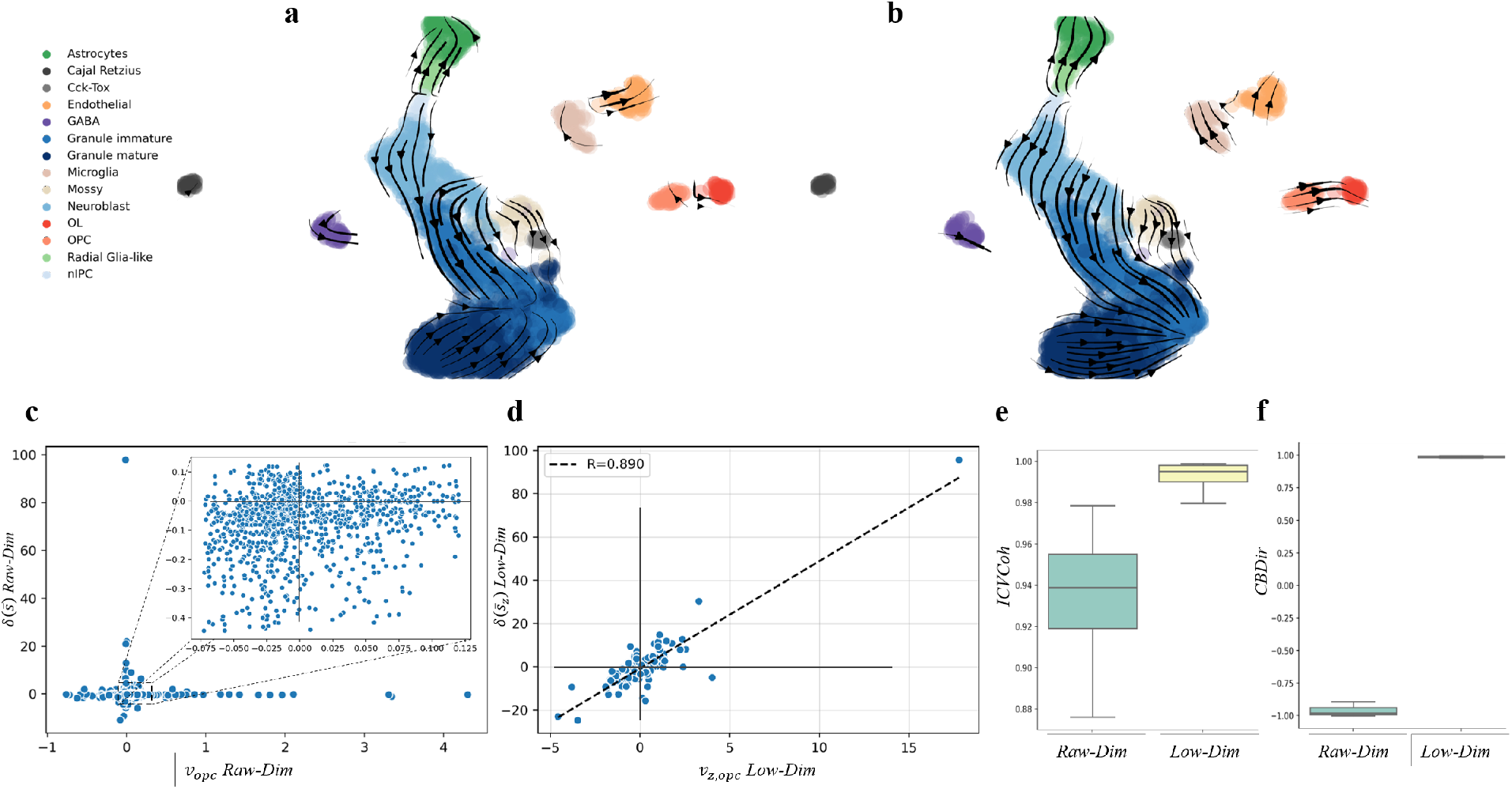
Results and Analysis on Dentategyrus. **a**, scVelo Stochastic Mode in Raw Gene Space; **b**, Velocity Projected into Low-Dimensional Space; **c**, Scatter plot of 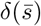 (OL - OPC) over 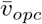 in the Raw Space; **d**, Scatter plot of 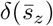 (OL-OPC) over 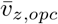 in the Low-Dim Space; **e**, In-cluster Coherence Scores; **f**, Cross-Boundary Direction Correctness (A->B) Scores.

However, we found the learned latent dimensions by veloAE show a strongly positive correlation between the velocity and expected differences (Pearsons’s R=0.890) and a high proportion of consistent signs (74.2%; Fig. 3d). Consequently, a clear direction from OPCs to OLs has been achieved by projecting on these lower dimensions (Fig. 3b), with increasing the direction correctness from −0.886 (scVelo stochastic) or −0.438 (scVelo dynamical) to 0.986 (veloAE; Fig. 3f and Table 1). Also, the transition directions are largely smoothed (in-cluster coherence: 0.992 by veloAE vs 0.936 by scVelo; Fig. 3e).

**Table 1:** Performance comparison across datasets between our proposed method veloAE, its variants and other baseline methods. Metrics include: in-cluster Coherence (ICVCoh) and Cross-boundary Direction Correctness (CBDir). The best performance in each metric is highlighted in bold font. Rankings of the veloAE are marked as superscripts. “w/” denotes with only the specific configuration. Stc: scVelo’s stochastic mode in raw space; Dyn: scVelo’s dynamical mode in raw space.

| Method | scNTseq |  | Dentate Gyrus |  | scEUseq |
| --- | --- | --- | --- | --- | --- |
|  | ICVCoh | CBDir | ICVCoh | CBDir | ICVCoh |
| scVelo (Stc) | 0.9141 | 0.2529 | 0.9363 | -0.8861 | 0.8297 |
| scVelo (Dyn) | 0.9527 | 0.2129 | 0.8204 | -0.4383 | 0.7517 |
| FA | 0.8289 | -0.1146 | 0.9454 | 0.0683 | 0.7945 |
| PCA | 0.9433 | 0.2758 | 0.9824 | -0.8126 | 0.9231 |
| AE | 0.9639 | 0.3143 | 0.9911 | -0.8164 | **0.9838** |
| veloAE | <sup>2</sup>0.9641 | <sup>2</sup>0.4024 | <sup>2</sup>0.9922 | <sup>1</sup>**0.9863** | <sup>3</sup>0.9709 |
| - AE w/ CohAgg | 0.9534 | **0.4055** | 0.9776 | 0.9473 | 0.9331 |
| - AE w/ AttComb | **0.9659** | -0.0958 | **0.9954** | 0.9846 | 0.9805 |

### 2.4 veloAE identifies intestinal organoid differentiation

In addition, we applied veloAE to an intestinal organoid data set, where a snapshot is taken during the differentiation from stem cells to secretory cells or enterocytes [17]. In the original paper, two strong differentiation trajectories have been identified by Monocle2 [3], and the directions were manually added by annotating the stem and differentiated cell types. Here, we ask whether RNA velocity can automatically identify the differentiation trajectory and its directionality. By using the spliced and unspliced RNAs, we found that the RNA velocity estimated by scVelo stochastic model is able to identify the branch 1 trajectory from stem cell to secretory cells, but fails to find the fully correct direction on the branch 2 terminating at enterocytes (Fig. 4a for stochastic model, Supp. Fig. S1 for dynamical mode). On this branch, two opposite directions were wrongly suggested by scVelo, possibly because the low cell density in the middle of this trajectory.

**Figure 4.**
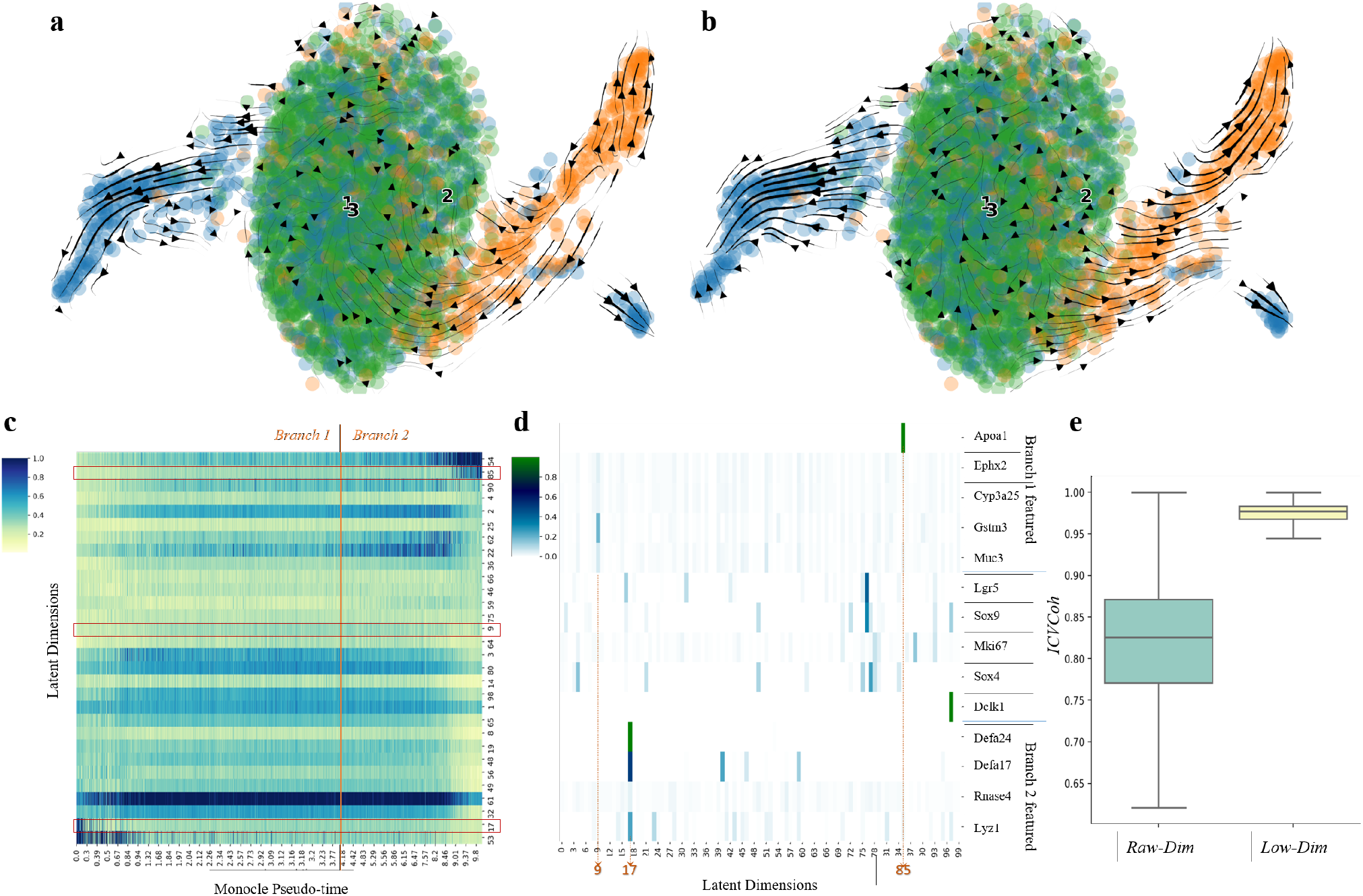
Results and Analysis on scEuseq. **a**, scVelo Stochastic Mode in Raw Gene Space; **b**, Velocity Projected into Low-Dimensional Space; **c**, Heat Map of Strongest 30 Low-Dim Expression Levels Along the Two Differentiation Branches (i.e., Monocle pseudo-time); **d**, Heat Map of Selected Genes’ Attention Distributions Over Latent Dimensions; **e**, In-cluster Coherence Scores.

By contrast, veloAE can successfully identify both differentiation trajectories and their directions through the learned lower dimensional representations (Fig. 4b). Interestingly, we found that the latent dimensions of expression states well display the dynamics of the two differentiation branches (top 30 dimensions in Fig. 4c). By examining the attention weights in the attentive combination module, we found that the major marker genes of the two differentiation trajectories dominantly enrich in three latent dimensions: branch 1 (secretory cells) genes Gstn3 in dim 9 and Apoa1 in dim 85 and branch 2 (enterocytes) genes Defa24, Defa17 Lyz1 in dim 17 (Fig. 4d). Furthermore, thanks to the effective representation, the estimated transition directions are significantly more coherent compared to the original gene space (0.830 by scVelo vs 0.971 by veloAE; Fig. 4e).

### 2.5 Comparison with multiple baseline methods

In order to assess the contribution of each module in our proposed method, we compared the velocity coherence and direction correctness between veloAE, and its two variants by keeping only the GCN in the encoder for cohort aggregation (i.e., w/ CohAgg) or only the attention module in the decoder (i.e., w/ AttComb), or by turning off both modules and changing it to a vanilla auto-encoder (AE). We also included principal component analysis (PCA) and factor analysis (FA) models as additional baseline methods.

Compared to scVelo, both PCA and all AE based methods consistently achieve higher within-cluster velocity coherence (ICVCoh) across all these three data sets (Table 1), probably thanks to their abilities of data denoise. Among them, veloAE remains comparable to the method with the highest ICVCoh in any data set. However, the ablation of the attention module in decoder (i.e., the variant: AE w/ CohAgg) results in obvious ICVCoh decreases, e.g., from 0.971 to 0.933 in scEUseq data set, suggesting the attentive combination is critical for smoothing the transition direction within a cell cluster.

Importantly, when examining the direction correctness, only veloAE and its variant with the GCN module for cohort aggregation (i.e., AE w/ CohAgg) can achieve high CBDir scores on both scNTseq and Dentate Gyrus data sets (Table 1). This highlights the importance of the cohort aggregation module for ensuring an informative projection of RNA velocities for trajectory inference. Taken together, veloAE by combining the GCN module in the encoder and attention module in the decoder, has a balanced performance on both high coherence and accurate directionality.

## 3 Discussion

In this work, we aimed to construct the cell transitions from RNA velocities, which is a new paradigm of trajectory inference that the intrinsic dynamics are implied from the two layers of molecular observations: the spliced and unsplcied RNAs. Computationally, cosine or correlation based metrics on vectors are used to measure the similarity between two cells in terms of directionality. As the velocity vectors are not self-normalized, such metrics are substantially different from Euclidean distance on transcription profiles that is commonly used in conventional trajectory inference methods. Therefore, a lower-dimensional representation with denser data could be crucial for the RNA velocity analysis, which motivates us to develop veloAE here, a tailored auto-encoder to learn effective representations for RNA velocity projection. Our findings from the evaluation experiments on three different datasets demonstrate the benefits of applying veloAE in post-processing raw velocity estimations for more robust results. The robustness of projections is impressively indicated in both visual demonstrations and quantitative metrics.

Our veloAE model consistently outperforms baseline methods across datasets thanks to the proposed two mechanisms. While *cohort aggregation* explicitly constrains a cell’s low-dimensional representation to resemble its neighboring cells, *attentive combination* urges the representations to retain as much information from the original gene space as possible for reconstructing the input. Compared with either PCA or a standard Autoencoder, VeloAE can hence naturally project similar cells closer to each other (effect of cohort aggregation), forming more centered neighborhood or clusters for later analysis. Meanwhile, by replacing the decoder (e.g., an MLP) in AE with the attentive combination mechanism, VeloAE can implicitly embed the similarity information among gene profiles into the low-dimensions, which may not be learned by the decoder of an AE instead. Consequently, the synergistic effect is that the low-dimensional cell representations can not only best keep the key information for reconstructing the original space but also explicitly encode its spatial closeness evidence among cells.

Finally, we would like to mention that the proposed method shares some merits with a bunch of dimensionality reduction methods in *manifold learning* [18], e.g., Isomap [19], Locally Linear Embedding (LLE) [20], Laplacian Eigenmap (LE) [21], t-distributed Stochastic Neighbor Embedding(t-SNE) [22], and Uniform Manifold Approximation and Projection (UMAP) [23]. Assuming high dimensional data to have concentrated low-dimensional intrinsic structures (i.e., manifolds), these methods attempt to map data from the original space into a low-dimensional space, where the main variation is kept along the manifold. Manifold learning methods usually rely heavily on the local structures of samples, as it is assumed that the local neighborhood of a sample can be approximated as a Euclidean space, which can be simply characterized by a k-nearest-neighborhood graph. Computed in the original space, such neighborhood graph participates in manifold learning and force the low-dimensional representation to retain the local structure of the original space. Our method, with the help of an encoder to concentrate information from raw data into low-dimensional space, also attempts to keep the local structures of both cells and genes by involving an explicit neighborhood graph (in cohort aggregation), and the implicit gene profile similarities (in attentive combination). That said, our approach shares the motivation with prior manifold learning methods, and we expect it could enlighten future applications of concepts from manifold learning in robust single cell velocity estimation.

## 4 Methods

### 4.1 RNA velocity Autoencoder (veloAE) model

A standard autoencoder consists of an encoder and a decoder [24] which are usually parameterized by multi-layer perceptrons (MLP) with a single hidden layer (See Supplementary Fig. S4). When processing data, an encoder transforms input data points into a low-dimensional space, where key information about the data is retained. A decoder then reconstructs the input dimensions from the dimension-reduced representations, aiming to recover the original input. The low-dimensional representations are of particular interests to us, as they, when preserving key information, are expected to filter noises or less important information, thus enabling more coherent estimations for single cell velocities.

Effective as it is, the standard autoencoding architecture is limited in incorporating inter-sample correlations into the encoding procedure. More specifically, a standard autoencoder assumes the input data to be independently identical distributed (IID), hence there is no explicit mechanism of keeping cells’ neighborhood information in the low-dimensional space. Yet it is unable to capture cross-cell distribution patterns of a gene to identify key latent dimensions for its reconstruction. It is, however, noted that the neighborhood information of cells plays a critical role in cell-level velocity inference, and that the steady-state approach of gene-level velocity inference also exploits cross-cell patterns of a gene.

To mitigate the above issues, this work proposes two mechanisms that can be embedded into the encoder and decoder modules of an autoencoder:

- Cohort aggregation with graph convolutional networks (GCN) [25] for incorporating neighborhood information into low-dimensional representations;
- Attentive combination of latent dimensions for target input dimension reconstruction.

The overall computational framework is illustrated in Supplementary Fig. S5. In the following text, we formalize our framework and the proposed two mechanisms.

Suppose an input matrix 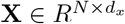denotes any matrix of transcriptome, spliced or unspliced reads, with normalized counts of *d*_*x*_ genes across N cells. The Encoder parameterized by a single-hidden layer MLP transforms it into a low-dimensional space as 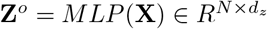, where *d*_*z*_ is much smaller than *d*_*x*_. To associate representations of cell neighborhood, the following cohort aggregation procedure is applied to **Z**^*o*^ with the help of a pre-specified neighborhood graph.

#### Cohort aggregation (CohAgg.)

leverages GCN to enrich the low-dimensional representation of a cell by taking the weighted average of its neighborhood including itself. Formally, an aggregated low-dimensional representation 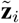 for the *i*th cell is computed as:

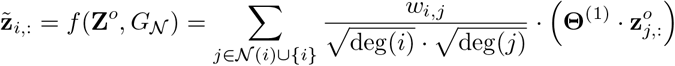

where *G*_*𝒩*_ is a given neighborhood graph (usually generated when scRNAseq data are processed), *deg*(*·*) returns node degree in the graph, *w*_*i,j*_ is the similarity score between cell i and j, and the set operation *𝒩* (*i*) *∪ {i}* groups cell *i* and its nearest neighbors in a common set. Θ^(*·*)^ is the learnable parameter matrix of GCN. In aggregation, *w*_*i,j*_ weighs the importance of neighbor representation *j*, and inverse of degrees normalizes the result.

Compactly, the computation can be conducted in matrix form as:

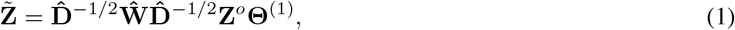

where 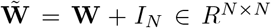is the weighted adjacency matrix **W** *∈ R*^*N×N*^ of the neighborhood graph *G*_*𝒩*_enhanced by self-loops of weight one, and 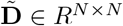 is the degree matrix with each diagonal element computed as 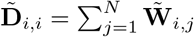.

Moreover, to encode the influence of second-order neighborhood, the same cohort aggregation mechanism is applied to 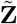, which yields the final encoding result **Z** which explicitly encodes cohort relationship into low-dimensional representations:

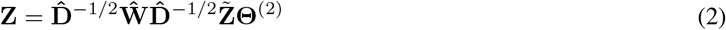

#### Attentive combination of latent dimensions (AttComb)

The latent dimensions of **Z** compress multiple original dimensions. Consequently, reconstructing an input dimension from **Z** requires querying information encoded in multiple latent columns. We adopt a global attention mechanism to evaluate the contribution of each latent dimension to a target input dimension.

To enable attention computation, a pre-specified representation matrix **G** for genes is required. The matrix offers in its each column 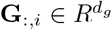a vectorized representation for a target gene. In practice, **G** can be prepared using techniques such as principle component analysis (PCA) on the columns of a count matrix, or learned from data.

Given a target gene represented as **G**_:,*i*_, reconstruction of its corresponding input dimension 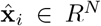 requires computing the attention weights for each of the latent dimensions in **Z**. To achieve this, we first transform the gene representation to a query vector 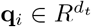, and each latent dimension **Z**_:,*j*_ to a key vector 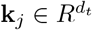, both with separately parameterized MLPs. Scaled dot product between each pair of the query and key vectors are then calculated and normalized with softmax function to yield the weights:

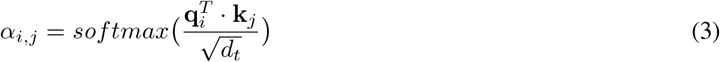

The weights are then used to aggregate corresponding dimensions of **Z** by weighted sum to reconstruct the target input dimension. For reconstructing the reads (denoted as 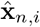 of the *i*th gene in the *n*th cell, the procedure works as follows:

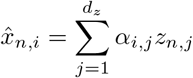

Compactly, the computation for reconstructing all the input dimensions can be conducted in matrix form as:

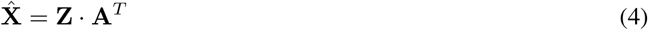

Note that the attention matrix **A** plays the role of a single fully-connected feedforward neural layer, mimicking a standard decoder without any hidden layer or nonlinearity. However, different from a standard decoder, each attention weight in **A** is computed based on the global information of both latent and target dimensions, instead of fitting locally under the IID assumption as a standard decoder does. Moreover, as a by product, the attention weights can help us better interpret the latent dimensions of **Z** by tracing their contributions to input dimensions.

#### Fitting and velocity projection

All the parameters of the framework are fitted using mean squared error (MSE) between reconstructed 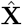 and the original **X**. To enable the model to simultaneously encode reads of transcriptome (**X**), spliced RNA (**S**) and unspliced RNA (**U**), we construct all the three matrices and add up their MSE losses to form the optimization objective:

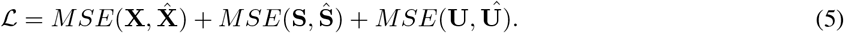

Note, **X** is often non-redundant to **S** and **U** due to their different methods for reads counting. Otherwise, it is optional to only use **S** and **U** to fit this model. After fitting the parameters, the encoder of the framework can project **X, S, U** into low-dimensional representations as **X**_*z*_ = *encode*(**X**), **S**_*z*_ = *encode*(**S**), **U**_*z*_ = *encode*(**U**), respectively. The velocity of cell i 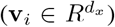 inferred in the input space can also be projected to the latent space using the formula: *encode*(**s**_*i*_ + **v**_*i*_) *− encode*(**s**_*i*_), which assumes that the mass difference of spliced RNA within a unit time span is the velocity in both input and low-dimensional space. Computationally, this treatment allows input to the encoder more similar to those at the training stage than directly projecting velocity vector. The compact form of projecting velocities of all cells using matrix algebra is:

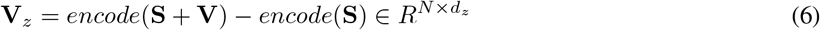

### 4.2 Cell transitions from low-dimensional representations

With low-dimensional projections **X**_*z*_, **U**_*z*_, **S**_*z*_, **V**_*z*_ ready, the estimation for cellular transitions can take place in the low-dimensional space and is expected to achieve more robust results. In accordance with scVelo, the transition strength *π*_*i,j*_ from cell *i* to cell *j* in the low-dimensional space is defined as the cosine similarity between the low-dimensional velocity of cell *i* and the difference of splicing expressions between cells *j* and *i*:

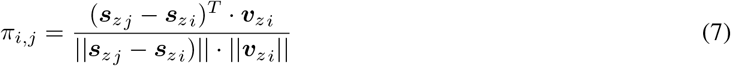

This transition score is then normalized with softmax function (with optional weights) over the neighborhood of cell *I* to yield the transition probabilities (details in [9]).

### 4.3 Settings of used models

To validate the performance of the proposed framework, evaluation experiments were conducted on multiple single-cell sequencing datasets, with baseline methods involved for comparison. On each dataset, we fit all the candidate models and estimate cell velocities using the scVelo package [9] in both the stochastic and dynamical modes. As scVelo uses the first-order moments (mean) of cell neighborhood instead of raw spliced and unspliced reads for estimation, to prevent mismatch we follow this practice and fit all the other candidate models (those in Sections 4.1 and 4.4) using the same moment matrices of spliced and unspliced reads.

The default data preparation procedures introduced in [9] and implemented in scVelo is adopted when we preprocess all the evaluation datasets: Top 2,000 highly variable genes passing a minimum threshold of 20 expressed counts are normalized and kept; A 30-neariest-neighbor graph is constructed based on 30 principle components in the PCA space for computing each cell’s first- and second-order expression moments, all using the scVelo methods:

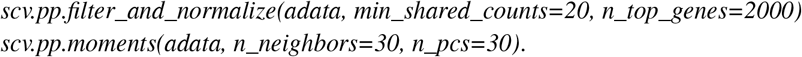

Then, regarding the stochastic mode, we apply the default settings in scVelo package and fit all the models with: *scv*.*tl*.*velocity(adata, mode=“stochastic”)*. Similarly, we adopt all the default configurations of the package to fit scVelo dynamical models, except that we increase the maximum iteration rounds from 10 to 100 for more sufficient fitting, and that we force all the 2000 genes involved in the estimation, which allows slight gains in quantitative performance metrics. The methods for fitting dynamical models are:

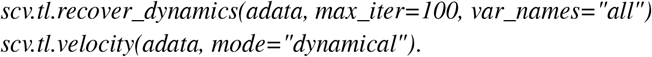

### 4.4 Baseline methods

All the hyper-parameters of the candidate models are summarized in Supplementary Table S2.

#### Factor Analysis (FA

[26] models the variance of observed variables with a potentially lower number of unobserved latent variables called factors. To make it comparable to our method, we fit FA on the concatenated matrices of transcriptome, spliced and unspliced expression reads, and use the same computation as Equation 6 to obtain the low-dimensional velocities.

#### Principle Component Analysis (PCA

[27] projects high-dimensional data into a low-dimensional space of orthogonal basis, while keeping the most variance of the original data. To make it comparable to the proposed framework, as FA we fit PCA on the concatenated matrices of transcriptome, spliced and unspliced reads, and use the same strategy as Equation 6 to obtain the dimension-reduced velocities.

#### Standard autoencode

It applies MLPs with single hidden layers to parameterize encoders and decoders, apart from which the same reconstruction task and projection strategy are adopted in fitting and applying the model.

#### Ablation: with only cohort aggregation (w/ CohAgg

In this ablation configuration, the attentive combination module is replaced with a single feedforward neural layer without any hidden layer or non-linearity. Such configured decoding layer is best comparable to the global attention weights in Equation 4, which allows us to compare the influence of attentive combination versus a standard decoder in input dimension reconstruction. This ablation model also shares the same reconstruction task and projection strategy with the proposed model.

#### Ablation: with only attentive combination (w/ AttCom

In this ablation configuration the GCN-enabled cohort aggregation module is removed, which is aimed to study the impact of incorporating neighborhood information in encoding the input dimensions. Similarly, the same reconstruction task and projection strategy is also adopted in this configuration.

#### Implementatio

We implemented the standard autoencoder, the proposed veloAE and the ablation models using pytorch. The GCN layers of veloAE and CohAgg are implemented using pytorch-geometric package. We choose GELU as the nonlinear activation for any MLP with hidden layer, and used AdamW for optimizing the model parameters. The specific learning rate is a key hyper-parameter decided seperately across datasets. Other hyper-parameters like the size of low-dimensional space *d*_*z*_ and number of epochs are set consistent over all relevant models and listed in Supplementary Table S2. PCA is implemented with scikit-learn package. All the neural network models are fit with a single NVIDIA GTX1080ti GPU.

### 4.5 Evaluation metrics

Prior works usually rely on visual observations on the plotted velocity estimation results to analyze and evaluate velocity estimation methods. However, we argue that visual judgment of velocity estimation results may be misleading and easily fall short in differentiating results that are less visually evident. Quantitative metrics hence should be developed. Unfortunately, the only off-the-shelf metric available is scVelo’s coherence score which indicates a cell’s general velocity coherence with its immediate and second-order neighbors.

Coherence alone, however, can not reliably demonstrate the correctness of velocity directions for cells with known differentiation order. To illustrate, it is very likely that the directions of velocities could be totally reversed against ground-truth differentiation orders, but still be coherent, thus retaining an unfair high score.

To allow for a more comprehensive evaluation, we propose below two types of metrics targeting direction correctness and velocity coherence respectively. Note that the metric with “cross-boundary” prefix require input of ground-truth development directions for pairs of cell clusters, e.g., A->B. The statistics would be computed for those boundary cells, i.e. cells of type A with type B in the neighborhood. Formally, We define boundary cells indexed by the source cells (type A), and represent the set as

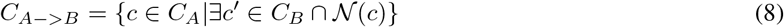

where *C*_*A*_ and *C*_*B*_ are sets of type A and B cells respectively, 𝒩 (*c*) retrieves the neighboring cells of c. The *∃* condition filters our non-boundary cells.

#### Cross-boundary Direction Correctness (A->B) (CBDir

is designed for evaluating how likely, following its current velocity, a cell can develop to a target cell. If a cell’s velocity is correctly estimated, then the velocity direction should be consistent with the development trajectory of the cell. Note that both source and target cells are represented in a common vector space, thus the development trajectory in a very short time period could be approximated simply with the location displacement from source to target cells in this space. Therefore, given a ground-truth developmental direction from cell type A to B, an ideal velocity of a type A cell is expected to be consistent with its displacement to a type B cell. Specifically, we consider only cross-boundary cells that mimic cell development at a very shot time, to prevent unexpected impacts of cell development within the same clusters, i.e., the development trajectory can hardly be approximated by simple displacement when a type B cell has developed to later stages greatly different from Type A.

The formula for computing this score is as:

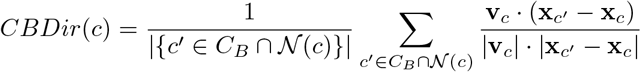

where **x**_*c*_′ and **x**_*c*_ are vectors representing cells c and c’ in a low-dimensional space via the Uniform Manifold Approximation and Projection (UMAP) algorithm [23], **x**_*c*_′ *−* **x**_*c*_ is the cell displacement in this space, and **v**_*c*_ is decomposed UMAP velocity representation in the same space. In the scVelo package UMAP representations can be computed using function: *scv*.*pl*.*velocity_embedding_stream*.

#### In-cluster Coherence (ICVCoh)

is computed with a cosine similarity scoring function between cell velocities within the same cluster. Specifically, the neighborhood of some cell c (e.g., of type A) is searched, and those cells within the same type (i.e., type A) are kept. Then, an average velocity similarity score for c and all these kept cells is computed as:

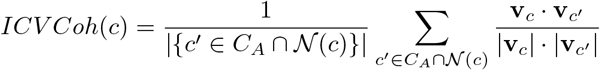

### 4.6 Datasets

To evaluate our methods, single cell sequencing datasets of three different biological systems are enrolled: scNTseq [10], scEUseq [17], and Dentate Gyrus neurogenesis [16]. There datasets contain known differentiation orders for all or part of the cell clusters, with which we can validate both the coherence and correctness of estimated velocities. The datasets’ information relevant to our experiment is summarized in Supplementary Table S1, and the ground-truth orders of cell development in the datasets are further elaborated below:

#### scNTseq

has time labels (in minutes) for each cell, which indicates cell development orders. We evaluate velocities between immediate consecutive time points as 0->15, 15->30, 30->60, and 60->120 respectively.

#### Dentate Gyrus

The known cell differentiation direction in Dentate Gyrus dataset is from oligodendrocyte precursor (OPC) to oligodendrocyte (OL) cells, which serves as the only ground truth for evaluation.

#### scEUse

groups all cells into three monocle branches. Although there are no inter-branch development orders known for applying our proposed metrics, cells within two branches (branches 1 and 2) exhibit uni-directional development trajectory. Therefore, correctly estimated velocities should be highly coherent within the two branching trajectories. We need only evaluate their in-branch coherence, and then check visually whether the velocites are pointing from the branching start to end to know their correctness.

### 4.7 Code availability

VeloAE is an open-source Python package available at https://github.com/qiaochen/VeloRep. All the analysis notebooks for reproducing the results are also available in this repository.

