## Supplementary File for "Representation learning of RNA velocity reveals robust cell transitions"

### Supplementary Materials for "Representation learning of RNA velocity reveals robust cell transitions"

Table S1: Summary of Datasets

| Dataset | # cells | # kept genes | known diff.order | eval.method |
|---|---|---|---|---|
| scNTseq [3] | 3,066 | 2,000 | labeled time order 0->15->30->60->120 | all |
| Dentate Gyrus neurogenesis [2] | 2,930 | 2,000 | OPC->OL | all |
| scEUseq [1] | 3,831 | 2,000 | cells within monocle branches 1 and 3 | in-cluster metrics |

Table S2: Summary of Model Hyper-parameters

| Model | $d_z$ | $d_t$ | MLP hidden size | # epochs | lr (scNT) | lr (scEU) | lr (dentategyrus) |
|---|---|---|---|---|---|---|---|
| VeloAE | 100 | 100 | 256 | 20,000 | $8.0 \times 10^{-7}$ | $1.0 \times 10^{-5}$ | $5.0 \times 10^{-6}$ |
| FA | 100 | – | – | – | – | – | – |
| PCA | 100 | – | – | – | – | – | – |
| Autoencoder | 100 | – | 256 | 20,000 | $1.0 \times 10^{-5}$ | $1.0 \times 10^{-5}$ | $1.0 \times 10^{-5}$ |
| AE w/ CohAgg | 100 | – | 256 | 20,000 | $5.0 \times 10^{-6}$ | $1.0 \times 10^{-6}$ | $1.0 \times 10^{-5}$ |
| AE w/ AttComb | 100 | 100 | 256 | 20,000 | $1.0 \times 10^{-6}$ | $3.0 \times 10^{-6}$ | $5.0 \times 10^{-6}$ |

**a.** scNTseq

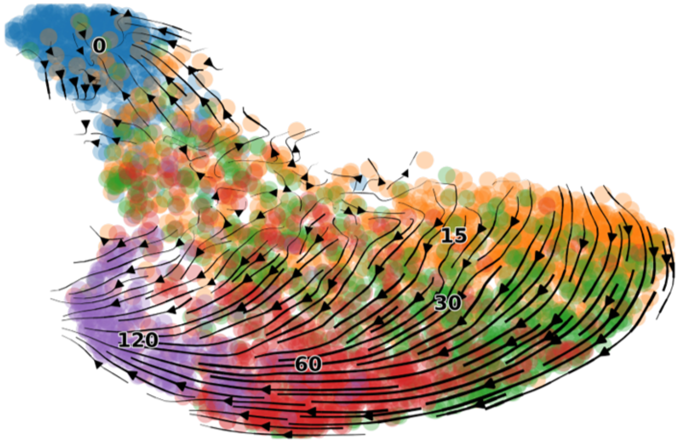


**b.** Dentate Gyrus Neurogenesis

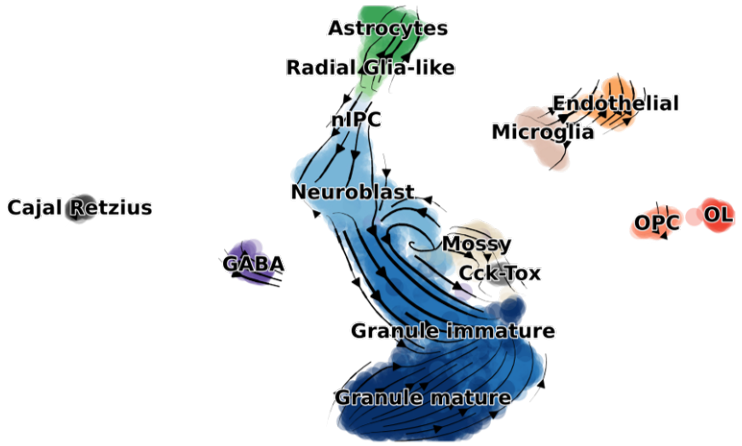


**c.** scEUseq

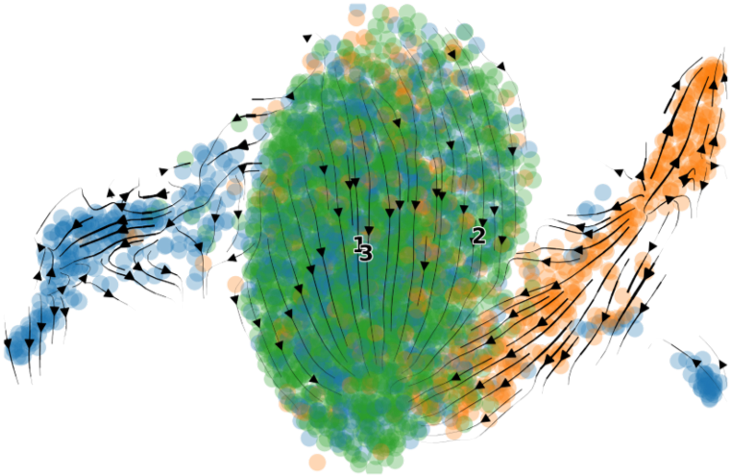


Figure S1: Velocity Estimated with scVelo Dynamical Mode

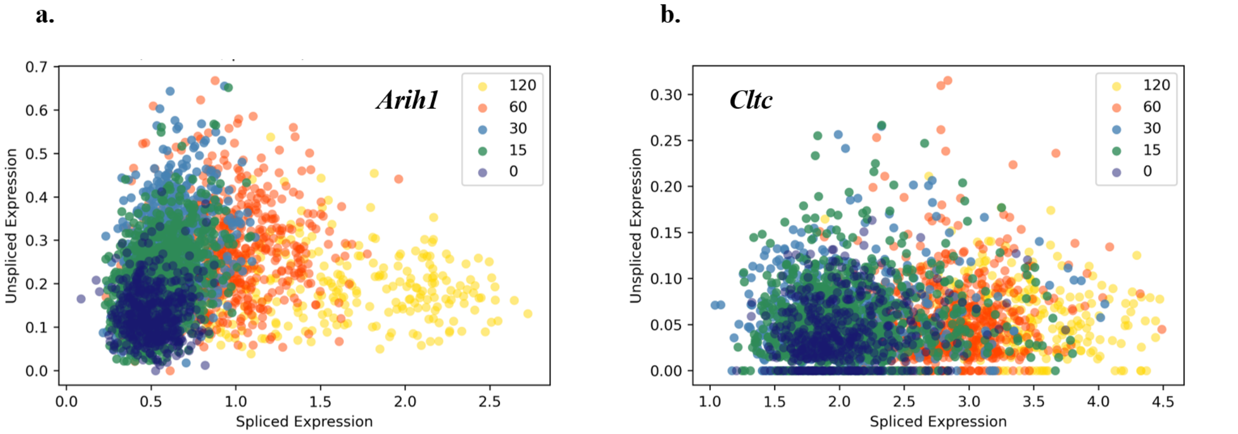


Figure S2: Plots of Example genes with Consistent (Arih1) and Inconsistent (Cltc) Velocity by Splicing Direction from 0 to 15 min on scNTseq. **a**. Plot of Spliced by Unspliced Expressions of Gene ***Arih1***. **b**. Plot of Spliced by Unspliced Expressions of Gene ***Cltc***.

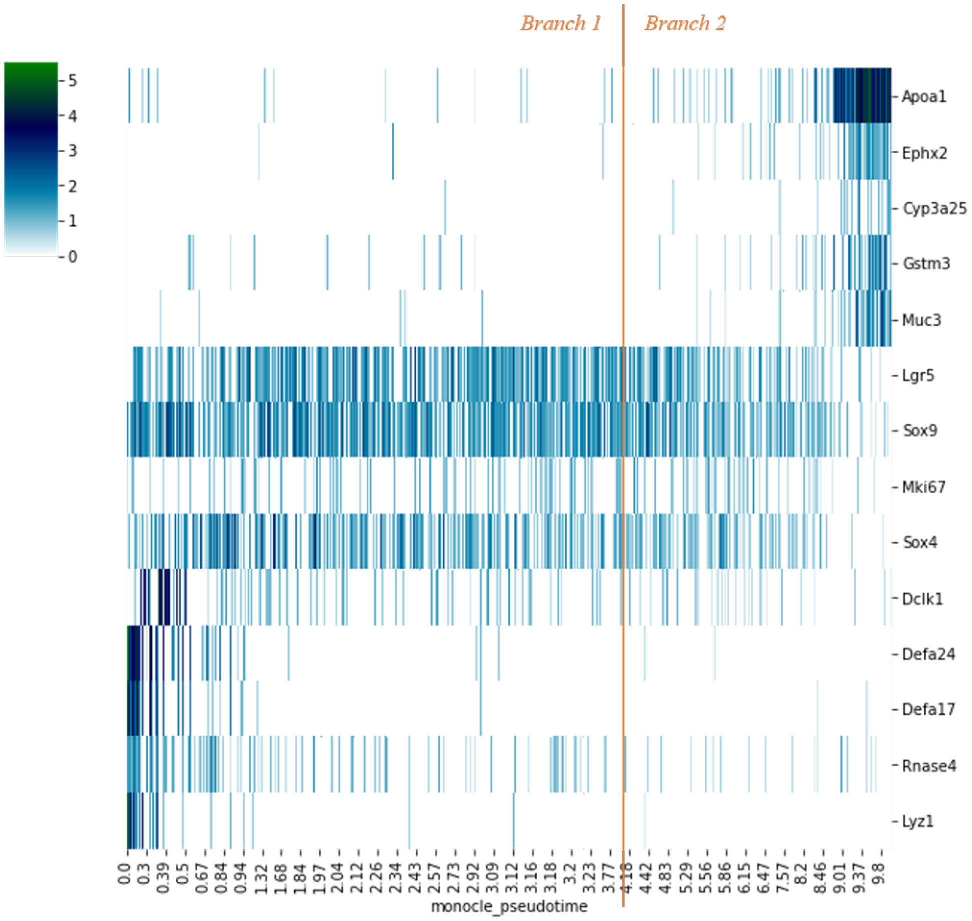


Figure S3: Expression Levels of Interesting Genes Over Two Branches (Ordered by Pseudo time).

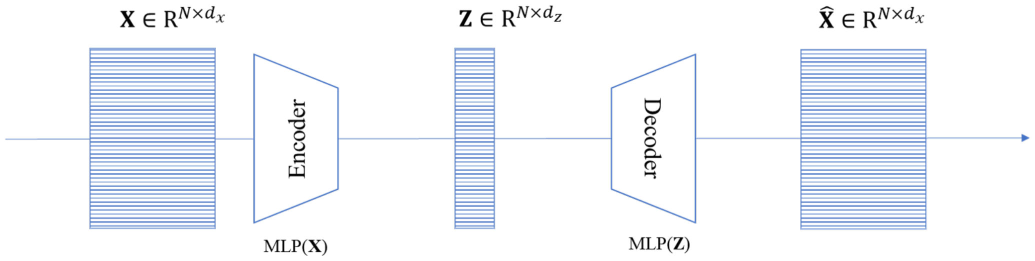


Figure S4: Architecture of a Standard Autoencoder

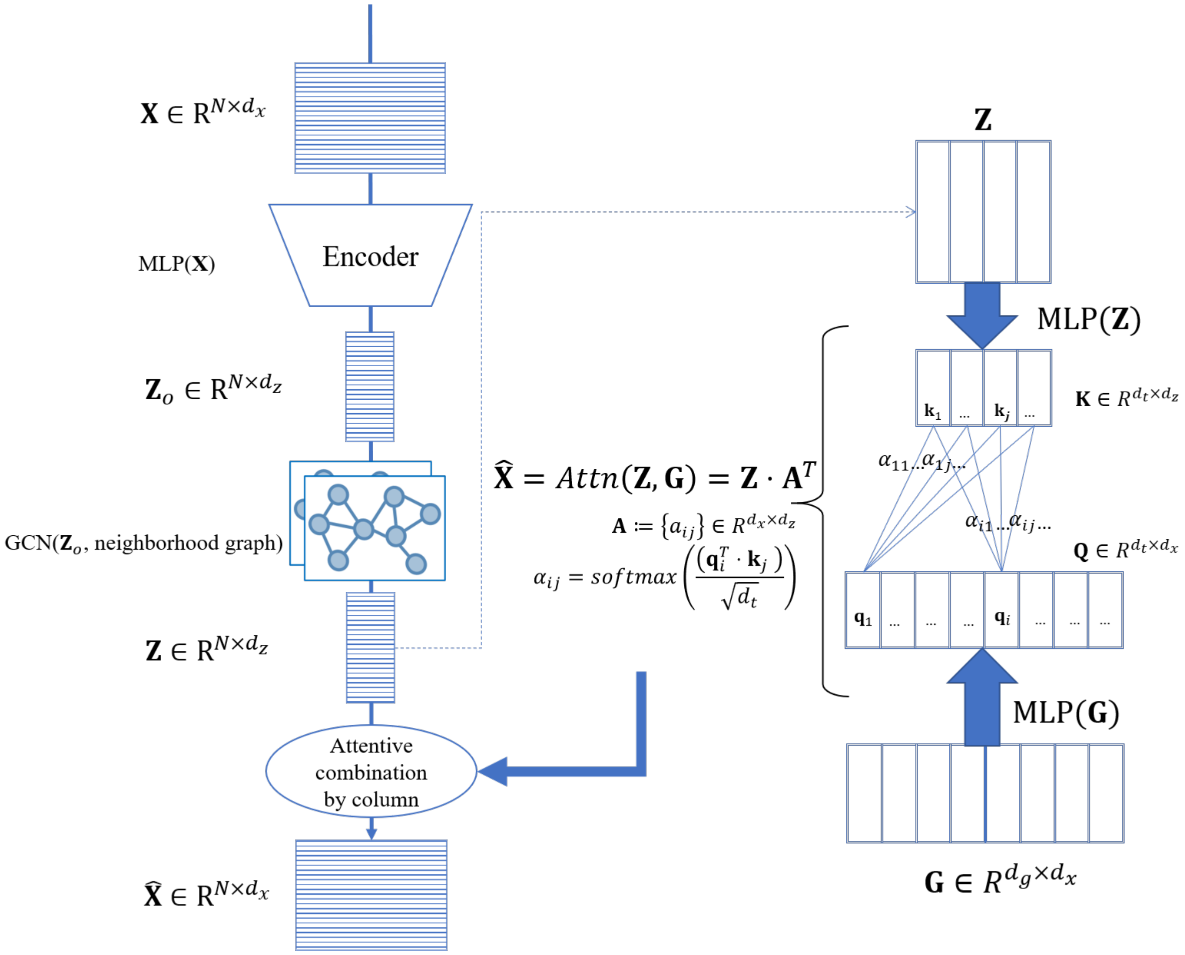


Figure S5: Architecture of the Proposed Computational Framework
